# Spatial Multi-Omics Reveals Extracellular Matrix Remodeling and VSMC Phenotypic Switching in Moyamoya Disease

**DOI:** 10.64898/2026.04.27.721225

**Authors:** Shihao He, Xiaofan Yu, Talha Ahmed, Yaoren Chang, Zhenyu Zhou, Hanzhi Liu, Yifan Xu, Junze Zhang, Shaoqi Xu, Lucas Du, Xun Ye, Rong Wang, Yuanli Zhao

## Abstract

**Background:** Moyamoya disease (MMD) is a progressive cerebrovascular disorder characterized by steno-occlusive lesions and intimal hyperplasia. Although vascular smooth muscle cell (VSMC) phenotypic switching is implicated in its pathogenesis, the precise spatial interplay between extracellular matrix (ECM) remodeling and local metabolic alterations within the distinct vascular microenvironments remains unknown.

**Methods:** Superficial temporal artery (STA) samples from patients with MMD and controls were analyzed by histology, immunofluorescence, spatial transcriptomics, spatial proteomics, and spatial metabolomics. Single cell RNA sequencing was used to profile the cellular landscape of STA tissues. To functionally validate the identified pathway, human brain vascular smooth muscle cells (HBVSMCs) were stimulated with fibronectin 1 (FN1), and patient derived induced pluripotent stem cell smooth muscle cells (iPSC-SMCs) were generated for migration and protein expression assays following ITGA5 silencing or focal adhesion kinase (FAK) inhibition.

**Results:** MMD STA samples exhibited marked intimal hyperplasia with medial thinning and intimal accumulation of α-SMA positive cells. Spatial transcriptomic and proteomic analyses identified an intimal remodeling program characterized by increased FN1, EFEMP1, fibronectin, ITGA5, and FAK, together with reduced MYH11. FN1 stimulation promoted smooth muscle cell migration, ECM associated protein expression, and FAK phosphorylation, whereas ITGA5 knockdown or FAK inhibition attenuated these effects. Patient derived MMD iPSC-SMCs showed similar abnormalities, including enhanced migration, increased FAK activation, reduced contractile markers, and increased ECM associated proteins. Spatial metabolomics and integrated multi-omics analyses further revealed that these changes were coupled to a metabolically depleted intimal niche enriched for reduced acyl-CoA related metabolites.

**Conclusions:** Integrated spatial multi-omics identifies coupled ECM remodeling and metabolic alteration in the hyperplastic intima of MMD. Within this context, the FN1-ITGA5-FAK axis emerges as a plausible mediator of smooth muscle remodeling that warrants further validation.

**Graphical Abstract:** 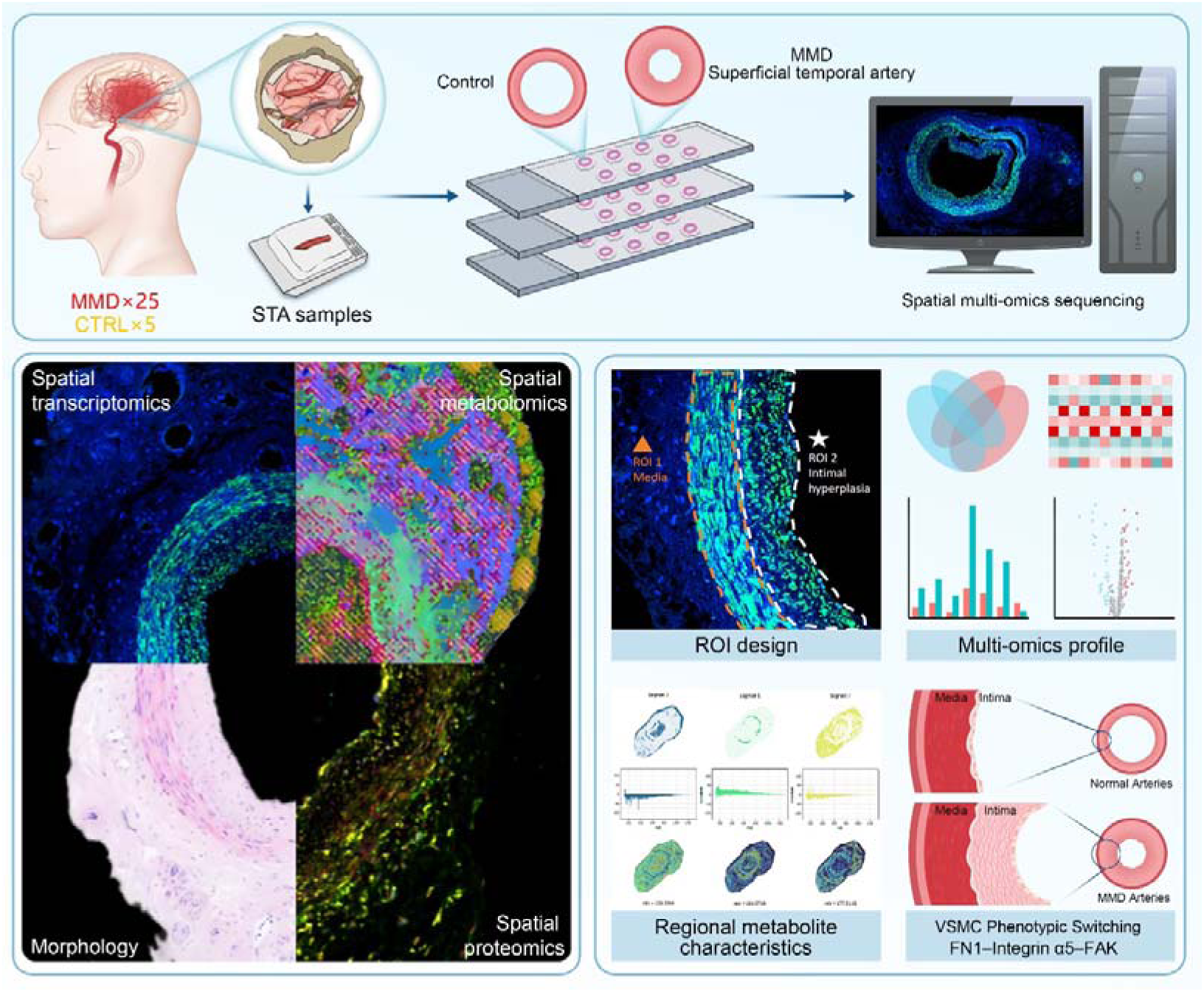

## Introduction

Moyamoya disease (MMD) is a chronic, progressive cerebrovascular occlusive arteriopathy, characterized by stenosis or occlusion of the terminal internal carotid arteries and their proximal branches, accompanied by the formation of an abnormal basal collateral circulation network.^1^MMD presents with diverse clinical manifestations, ranging from transient ischemic attacks to life-threatening intracranial hemorrhage.^2^ Despite decades of clinical observation, the precise pathogenesis of MMD remains largely elusive. While genetic predisposition, particularly the *RNF213* variant (e.g., p.R4810K), has been identified as a major susceptibility factor, genetic mutations alone cannot fully explain the complex spatiotemporal progression of the vascular lesions, suggesting a critical but unexplored role for local microenvironmental factors in driving the disease.^3^

The histological features of intracranial arteries in MMD involve pronounced intimal hyperplasia, thinning of the tunica media, and progressive waving or disruption of the internal elastic lamina.^4^ Classical hematoxylin and eosin (H&E) and immunofluorescence (IF) staining of staining of affected vessels (such as the middle cerebral artery and superficial temporal artery) both suggest that the luminal stenosis of arteries in Moyamoya disease may be caused by intimal hyperplasia.^5^ Unlike atherosclerotic plaques, which are primarily driven by lipid deposition and foam cell formation, the thickened intima in MMD is mainly composed of cellular components such as smooth muscle cells, leading to abnormal vascular remodeling and hemodynamic changes.^6^ Masuda et al. suggested proliferating smooth muscle cells within occlusive intracranial arterial lesions together with macrophage and T-cell localization, supporting the view that intimal expansion is an active cellular process rather than a purely degenerative change.^7^ Subsequent immunopathological analyses further showed α-SMA–positive cells in the thickened intima, vacuolar degeneration of medial smooth muscle cells, and strong S100A4 and IgG signals in the affected vascular wall, implicating phenotypically altered VSMCs and immune-related injury in the remodeling process.^8^ Similar intimal changes have also been documented in superficial temporal arteries (STAs) from patients with MMD, suggesting that the disease is not restricted to intracranial arteries alone and that surgically accessible extracranial arteries may capture relevant aspects of the vasculopathy.^9,10^ Considerable efforts have been made to elucidate the cellular origins and molecular mechanisms underlying this intimal hyperplasia. Previous *in vitro* and bulk-tissue studies have proposed that endothelial cell dysfunction, aberrant angiogenesis, and immune cell infiltration contribute to MMD pathogenesis.^11,12^ Importantly, vascular smooth muscle cells (VSMCs) are increasingly recognized as the central effectors in this occlusive process. Emerging evidence indicates that under chronic stress or genetic disruption, VSMCs undergo a profound phenotypic switching from a quiescent, contractile state to a highly proliferative, migratory, and synthetic state.^13^ Previous batch RNA sequencing or standard single-cell transcriptomics methods have been unable to preserve the spatial structure of tissues.

Although our previous single-cell transcriptomics results suggested that there may be an increase in smooth muscle cells in the vascular control group of Moyamoya disease^14^, there is still an important gap: the specific mechanisms of spatial changes in the metabolic microenvironment and VSMC in different pathological microenvironments (media and intima) are entirely unknown.

Recently, spatial multi-omics technologies, including spatial transcriptomics and mass spectrometry-based spatial metabolomics, have revolutionized our understanding of vascular diseases by preserving tissue architecture while providing high-throughput molecular mapping. In atherosclerosis, spatial transcriptomics has successfully localized specific macrophage-VSMC interactions and spatially confined inflammatory responses within the plaque microenvironment.^15^ Likewise, in abdominal aortic aneurysm and human carotid atherosclerosis, integrated single-cell and spatial transcriptomic approaches have revealed spatially organized stromal, immune, and VSMC states that shape disease progression and tissue instability.^16^ However, the spatially restricted metabolic and transcriptional programs that drive localized intimal hyperplasia remain undefined in moyamoya disease.

To bridge this gap, we mapped the spatial multi-omics atlas of superficial temporal arteries in MMD. By comparing the proliferating intima and thinning media, we discovered a robust extracellular matrix (ECM) remodeling program driven by the FN1-integrin α5-FAK axis, along with corresponding alterations in the local metabolic environment. Furthermore, we validated in vitro using a patient-derived induced pluripotent stem cell (iPSC) model that this signaling pathway may be involved in the pathological phenotypic transformation of MMD vascular smooth muscle cells (VSMCs).

## Methods

### Study participants

Patients examined between August 2024 and November 2025 with MMD were clinically diagnosed by medical history and neuroradiological examination, including cerebral digital subtraction angiography, magnetic resonance imaging, and functional regional cerebral blood flow studies following the diagnostic guidelines.^17^ The clinical data of all the patients were recorded. Clinical and imaging data were obtained from medical records and Picture Archiving and Communication Systems, respectively. Arterial segments (1–3 mm) were surgically excised from the STA of patients with MMD during revascularization (direct or combined bypass). For the control group, STA samples were obtained from patients undergoing craniotomy for epilepsy management. Written informed consent was obtained from all participants. This study was approved by the Institutional Ethics Committee of Peking Union Medical College Hospital, Beijing, China (I-24YSB0160).

### Histological and Immunofluorescence Analysis

Hematoxylin and Eosin (H&E) StainingTo evaluate the morphological structure of the superficial temporal artery (STA), 5 μm thick FFPE sections were subjected to H&E staining. Briefly, the sections were deparaffinized in xylene and rehydrated through a series of graded ethanol washes. After rinsing with deionized water, the slides were stained with Harris hematoxylin solution for 5 minutes, followed by differentiation in 1% acid alcohol. The sections were then counterstained with eosin Y solution for 2 minutes. Finally, the slides were dehydrated, cleared in xylene, and mounted with synthetic resin for light microscopy. The degree of intimal hyperplasia was quantified as the ratio of the intimal area to the medial area using ImageJ/Fiji software.

### Immunofluorescence (IF) Staining

Immunofluorescence was performed to visualize the expression and spatial distribution of vascular smooth muscle cell (VSMC) markers and integrins. FFPE sections were deparaffinized and rehydrated as described above. Heat-induced antigen retrieval was performed in Tris-EDTA buffer (pH 9.0) at 95°C for 20 minutes. To minimize non-specific binding, sections were blocked with 5% bovine serum albumin (BSA) in PBS at room temperature for 1 hour. The sections were then incubated overnight at 4°C with the following primary antibodies: anti-alpha Smooth Muscle Actin (alpha-SMA) (ab5694, Abcam; 1:200 dilution), anti-Integrin alpha 5 (ITGA5) (A19069, ABclonal; 1:200 dilution). After washing with PBS, the slides were incubated with fluorophore-conjugated secondary antibodies (Alexa Fluor 488 or 594, 1:500; Thermo Fisher) for 1 hour at room temperature in the dark. Nuclei were counterstained with DAPI (1:1000).

### Spatial Transcriptomics and Proteomics using GeoMx Digital Spatial Profiling (DSP)

Tissue Preparation and Probe Hybridization 5 μm thick formalin-fixed paraffin-embedded (FFPE) tissue sections were deparaffinized and subjected to target antigen retrieval using Tris-EDTA buffer (pH 9.0; Thermo Fisher). RNA targets were exposed via Proteinase K digestion (Thermo Fisher) and subsequently post-fixed in 10% neutral buffered formalin. The slides were incubated overnight at 37°C in a hybridization chamber with GeoMx Whole-Transcriptome Atlas (WTA) probes. To remove off-target probes, stringent washes were performed using saline-sodium citrate (SSC; Sigma-Aldrich) and 100% deionized formamide (VWR).

Morphology Staining and Region of Interest (ROI) Collection Following hybridization, tissues were stained with a fluorescent morphology marker panel to delineate distinct cellular compartments: Anti-alpha smooth muscle Actin antibody for vascular smooth muscle cells (1:500; Abcam, ab5694). Slides were loaded into the GeoMx DSP instrument (NanoString) for immunofluorescence imaging. Based on the morphological marking of the artery, the ROI is selected and segmented into marker-specific areas of illumination (AOIs), such as the media and intima, followed by spatially indexed UV-mediated barcode cleavage and collection.

Library Preparation and Sequencing Collected barcodes were indexed and amplified using the manufacturer-supplied PCR master mix. The resulting amplicons were pooled and purified via AMPure XP beads and ethanol washes. Library quality and size distribution were assessed using a Bioanalyzer high-sensitivity DNA trace, and the final libraries were sequenced on an Illumina NovaSeq 6000 platform.

Data Processing and Bioinformatics Analysis Raw FASTQ files were processed into deduplicated count matrices based on unique molecular identifiers (UMIs) and target tags. The limit of quantitation (LOQ) was established as the geometric mean plus two geometric standard deviations of the negative control probes. Genes consistently falling below the LOQ were removed, and the filtered datasets were normalized using upper quartile (Q3) normalization.

For differential expression analysis, counts were transformed to a log2 scale and normalized to library size using the rlog function. Differentially expressed genes (DEGs) between groups were identified using the edgeR package. Statistical significance was defined as a false discovery rate (FDR) adjusted P-value < 0.05 and an absolute log2 fold change (|LFC|) ≥ 0.5. Pathway enrichment analysis of the DEGs was conducted using the ClusterProfiler R package, with P-values adjusted via the Benjamini-Hochberg correction. Finally, spatial cell-type deconvolution was performed using the SpatialDecon algorithm based on a custom reference cell profile matrix.

### Spatial Metabolomics via MALDI-timsTOF Imaging Sample Preparation

Serial cryosections (adjacent to those used for spatial transcriptomics and proteomics) were prepared to ensure precise spatial correlation across multi-omics layers. 5 μm Sections were mounted onto indium tin oxide (ITO)-coated glass slides and desiccated prior to matrix application.

### Matrix Coating

Matrix application was performed using an automated vibration-spray system. The sections were coated with 2,5-dihydroxybenzoic acid (DHB, 15 mg/mL) dissolved in an acetonitrile/water mixture (90:10, v/v). The spraying parameters were optimized for high-resolution imaging: a nozzle temperature of 60°C, a flow rate of 0.12 mL/min, and a nitrogen gas pressure of 10 psi. A total of 30 passes were applied to the slides, with a 5-second drying interval between successive passes to ensure a uniform and fine-grained matrix crystal layer.

### Mass Spectrometry Imaging (MSI) Acquisition

MALDI-timsTOF MSI data were acquired using a timsTOF FleX system (Bruker Daltonics, Bremen, Germany) equipped with a 10 kHz SmartBeam 3D laser. The laser power was optimized at 80% and maintained constant throughout the session to ensure intra-sample consistency. Spectra were recorded in positive ion mode across a mass-to-charge ratio (m/z) range of 50–1300 Da. For high-resolution spatial mapping, the laser spot size (spatial resolution) was set to 5 \mum, with each pixel consisting of 400 accumulated laser shots.

Data Processing and Metabolite Identification

The raw MALDI mass spectra were normalized using the Root Mean Square (RMS) method to minimize ion-source variability, and signal intensities were visualized as normalized ion abundance maps. For definitive structural annotation of the identified metabolites, on-tissue MS/MS fragmentation was performed in dDA (data-dependent acquisition) mode on the timsTOF FleX system, allowing for detailed structural confirmation of key metabolic markers.

### Single-cell RNA sequencing and cell annotation

Superficial temporal artery (STA) samples were collected from 2 patients with moyamoya disease and 3 control subjects for single-cell RNA sequencing. Single-cell suspensions were processed using the 10x Genomics Chromium platform to generate 3′ gene expression libraries, followed by sequencing on an Illumina NovaSeq 6000 platform. After quality control, dimensionality reduction, clustering, and batch correction, cells were annotated according to canonical marker genes of major vascular and immune cell populations. Detailed experimental and bioinformatic procedures were the same as those described previously.^14^

### Generation and Maintenance of Patient-Derived iPSCs Isolation of Peripheral Blood Mononuclear Cells (PBMCs)

To establish patient-specific induced pluripotent stem cell (iPSC) lines, approximately 8–10 mL of peripheral venous blood was collected from MMD patients and healthy controls into anticoagulant tubes. Peripheral blood mononuclear cells (PBMCs) were isolated via density gradient centrifugation using Ficoll-Paque medium. Briefly, the diluted blood was layered onto the separation medium and centrifuged at 800 × g for 20–30 minutes at room temperature. The resulting PBMC layer was harvested, washed with dilution buffer, and resuspended in complete PBMC culture medium.

### iPSC Reprogramming and Colony Selection

The isolated PBMCs were reprogrammed into iPSCs using the non-integrating Sendai virus-based CytoTune™ 2.0 iPS Reprogramming Kit (Thermo Fisher Scientific) according to the manufacturer’s instructions. On day 0, cells were transduced with the Sendai reprogramming vectors and incubated overnight. Starting from day 3, the transduced cells were transferred onto rhVTN-N coated dishes and maintained in StemPro-34 medium. The culture was gradually transitioned to complete iPSC maintenance medium (e.g., mTeSR1) between days 7 and 8, with cells subsequently cultured on Matrigel-coated plates. Emerging iPSC colonies were monitored daily and identified based on characteristic pluripotent morphology. High-quality, undifferentiated colonies were manually picked, expanded, and purified for further characterization.

### iPSC Maintenance and Characterization

Established iPSC lines were maintained on Matrigel-coated dishes in mTeSR1 medium at 37°C in a humidified atmosphere with 5% CO_2_. The medium was replaced daily, and cells were passaged every 4–6 days using EDTA or Accutase upon reaching 80–90% confluency. The pluripotency of the iPSC lines was validated through immunofluorescence staining for markers such as OCT4, SOX2, and NANOG, as well as their capacity for tri-lineage differentiation (data not shown).

### Differentiation of iPSCs into vascular smooth muscle cells

Human iPSCs were dissociated into single cells and seeded onto Matrigel-coated culture plates at a density of 1 × 10^4^ cells/cm² in mTeSR1 medium supplemented with Y-27632. Mesoderm induction was initiated by replacing the medium with N2B27 containing CHIR99021 and BMP4 for 3 days. The cells were then switched to N2B27 supplemented with PDGF-BB and Activin A to promote smooth muscle lineage commitment, followed by further maturation in N2B27 containing Activin A and heparin. After day 9, the differentiated cells were maintained in smooth muscle cell medium for expansion and downstream experiments. For routine culture after differentiation or recovery from cryopreservation, cells were plated on Matrigel-coated plates and cultured in smooth muscle cell medium (ScienCell, #1101) under standard conditions (37 °C, 5% CO2), with daily medium changes.

### Human brain vascular smooth muscle cells culture

uman brain vascular smooth muscle cells (HBVSMCs; ScienCell, 1106) were cultured in smooth muscle cell medium (SMCM, ScienCell, 1101) supplemented with 2% fetal bovine serum, 1% smooth muscle cell growth supplement, and 1% penicillin–streptomycin.

HC-iPSC-SMCs and MMD-iPSC-SMCs provided by the authors were maintained in smooth muscle cell-specific medium. Cells were incubated at 37 °C in a humidified atmosphere containing 5% CO2. When cell confluence reached ∼90%, cells were washed twice with PBS and dissociated with 0.25% trypsin–0.02% EDTA. Digestion was terminated with complete medium, and cells were collected by centrifugation and passaged for subsequent experiments.

In the first, HBVSMCs were assigned to the following groups: control, 10 μg/mL FN1, 10 μg/mL FN1 + shRNA-NC, 10 μg/mL FN1 + shRNA-ITGA5, and 10 μg/mL FN1 + 1 μM VS6063. For FN1 stimulation, culture plates were pre-coated with human FN1 at 10 μg/mL, and cells were incubated for 24 h. In the second setting, HC-iPSC-SMCs and MMD-iPSC-SMCs were divided into HC-iPSC-SMC, MMD-iPSC-SMC, MMD-iPSC-SMC + shRNA-NC, MMD-iPSC-SMC + shRNA-ITGA5, and MMD-iPSC-SMC + 1 μM VS6063 groups. After treatment for 24 h, cell migration and protein expression analyses were performed.

### Lentiviral transduction

For ITGA5 knockdown, cells were transduced with lentiviral vectors carrying shRNA-ITGA5 or control shRNA-NC. One day before infection, cells were seeded in complete medium. Viral transduction was performed in the presence of infection reagent according to the indicated culture vessel volume, and cells were incubated for 12 h before replacement with fresh medium. Cells were then cultured for 3 additional days. Knockdown efficiency was validated by qPCR and western blotting before downstream experiments.

### Quantitative real-time PCR

Total RNA was extracted from treated cells using TRIzol reagent according to the manufacturer’s instructions. Briefly, cells were lysed in TRIzol, followed by chloroform extraction and isopropanol precipitation. RNA pellets were washed with 75% ethanol, air-dried, and dissolved in DEPC-treated water. RNA concentration and purity were measured using a NanoDrop 2000 spectrophotometer. cDNA was synthesized from total RNA using the iScript cDNA synthesis kit. Quantitative PCR was performed using TB Green Premix Ex Taq on an ABI 7500 real-time PCR system. Relative gene expression was calculated using the 2^-ΔΔCt method with GAPDH as the internal control. Primer sequences for ITGA5 and GAPDH are listed in the Table S1.

### Transwell migration assay

Migration of HC-iPSC-SMCs and MMD-iPSC-SMCs was evaluated using Transwell chambers. After treatment, cells were harvested, resuspended in serum-free medium, and adjusted to 2.5 × 10^5^ cells/mL. A total of 100 μL cell suspension was added to the upper chamber, while 600 μL DMEM containing 10% FBS was added to the lower chamber as a chemoattractant. After incubation for 24 h at 37 °C, cells on the upper surface were removed gently with a cotton swab. Cells that had migrated to the lower surface were fixed with methanol, stained with 1% crystal violet, and counted under a microscope.

### Western blotting

Total protein was extracted from treated cells using RIPA lysis buffer supplemented with PMSF. Protein concentrations were determined using a BCA protein assay kit. Equal amounts of protein (60 μg per sample) were separated by SDS-PAGE and transferred onto PVDF membranes. Membranes were blocked with 5% nonfat milk in TBST for 1 h at room temperature and then incubated overnight at 4 °C with primary antibodies against MYH11, CNN1, TAGLN, FN1, COL1A1, COL3A1, OGN, EFEMP1, ITGA5, FAK, p-FAK (Tyr397), and GAPDH. After washing, membranes were incubated with HRP-conjugated secondary antibodies for 1 h at room temperature. Protein bands were detected using enhanced chemiluminescence and visualized on a Tanon 5200 imaging system. Band intensities were quantified with Image Pro Plus 6.0 and normalized to GAPDH. Antibody dilutions are detailed in the report, including 1:1000 for most target proteins, 1:5000 for CNN1, and 1:10000 for GAPDH and secondary antibodies.

### Statistical analysis

All data are presented as mean ± SD from three independent experiments. Statistical analyses were performed using GraphPad Prism 9, and figures were assembled in Adobe Illustrator 2022. Differences among multiple groups were analyzed by one-way ANOVA followed by Tukey’s multiple-comparisons test. A P value < 0.05 was considered statistically significant.

## Results

### Histopathological characterization

To characterize vascular remodeling in MMD, we collected STA samples from 25 patients with MMD and 5 control subjects and performed histological staining, immunofluorescence, and spatial multi-omics profiling, including spatial transcriptomics, spatial proteomics, and spatial metabolomics (Fig. 1A).

**Figure 1.**
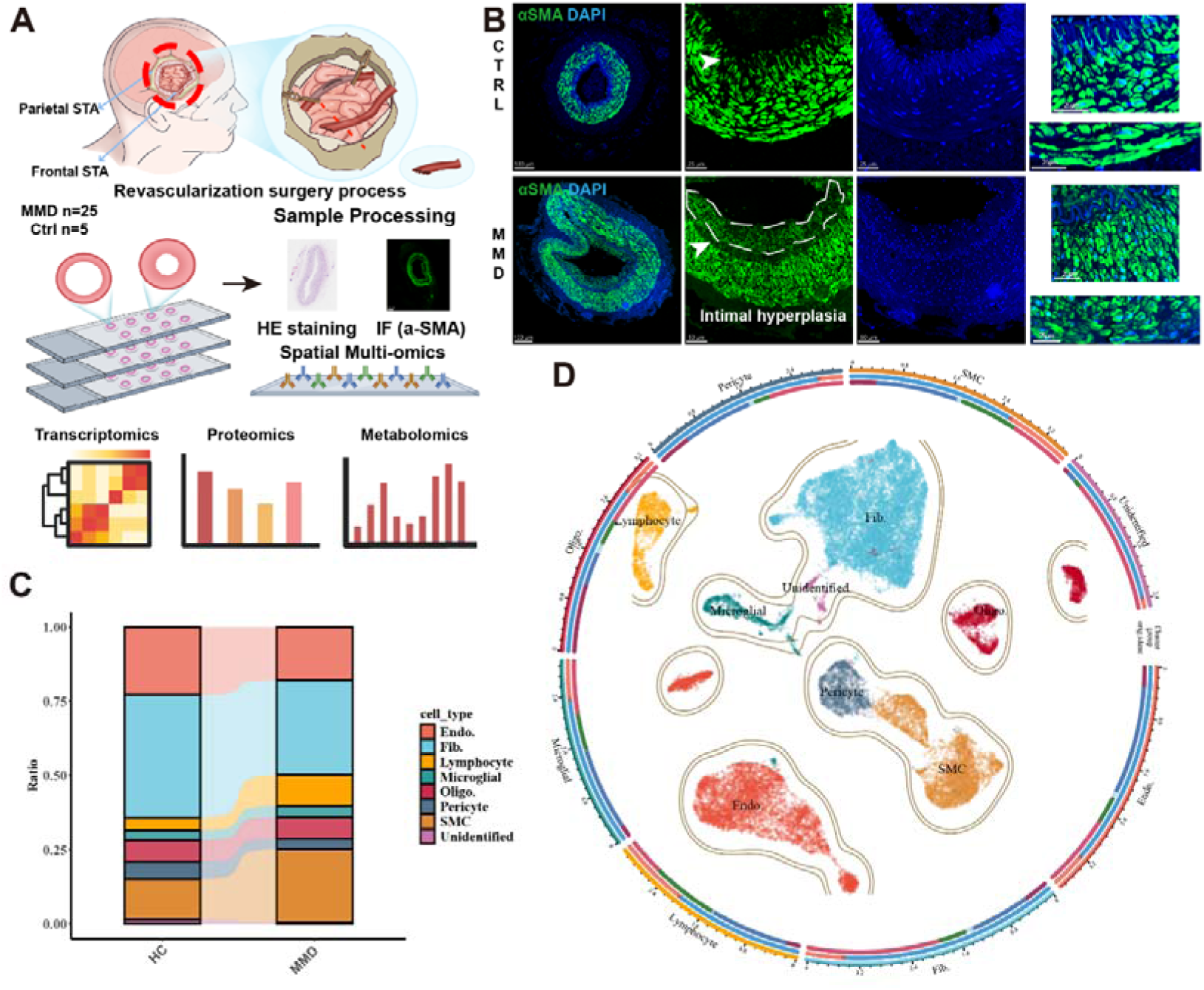
Spatial multi-omics landscape of Moyamoya disease arteries reveals SMC-dominated intimal hyperplasia. (A) Schematic overview of the experimental design. Superficial temporal arteries (STA) from Moyamoya disease (MMD) patients and controls were subjected to spatial transcriptomics, proteomics, and metabolomics. (B) Representative immunofluorescence (IF) staining of α-SMA (green) and DAPI (blue) in control and MMD arteries, highlighting severe intimal hyperplasia in MMD (white dotted lines). (C) Bar plot showing the relative ratio of distinct cell populations in healthy control (HC) and MMD vessels based on deconvolution analysis, indicating a significant expansion of the SMC compartment in MMD. (D) Circular plot visualizing the spatial clustering and cellular heterogeneity, mapping the distribution of SMCs, fibroblasts (Fib.), endothelial cells (Endo.), and local immune cells within the vascular microenvironment.

Compared with control vessels, MMD arteries exhibited marked intimal hyperplasia and a narrowed lumen, accompanied by abnormal accumulation of α-SMA-positive cells within the thickened intimal region (Fig. 1B). These findings are consistent with smooth muscle cell-associated vascular remodeling and support the presence of active intimal expansion in MMD. To further define the cellular landscape underlying these pathological changes, we performed single-cell analysis and identified multiple major cell populations in STA tissues, including endothelial cells, fibroblasts, lymphocytes, microglia, oligodendrocyte-like cells, pericytes, smooth muscle cells, and a small unidentified cluster. Comparative analysis of cell composition revealed a shift in the relative abundance of several cell populations in MMD compared with controls, indicating substantial remodeling of the vascular microenvironment (Fig. 1C, D).

### The hyperplastic intima of MMD STA exhibits ECM remodeling and activation of the ITGA5–FAK axis

To define the molecular features of intimal hyperplasia in moyamoya disease (MMD), we first compared the intimal and medial regions in superficial temporal artery (STA) sections based on histological annotation. Consistent with the histopathological findings, MMD samples showed marked intimal thickening accompanied by medial thinning, whereas control vessels displayed a relatively preserved vascular architecture (Fig. 2A). Quantification of the intima-to-media area ratio further confirmed a significant increase in intimal hyperplasia in the MMD group compared with controls (Fig. 2D).

**Figure 2.**
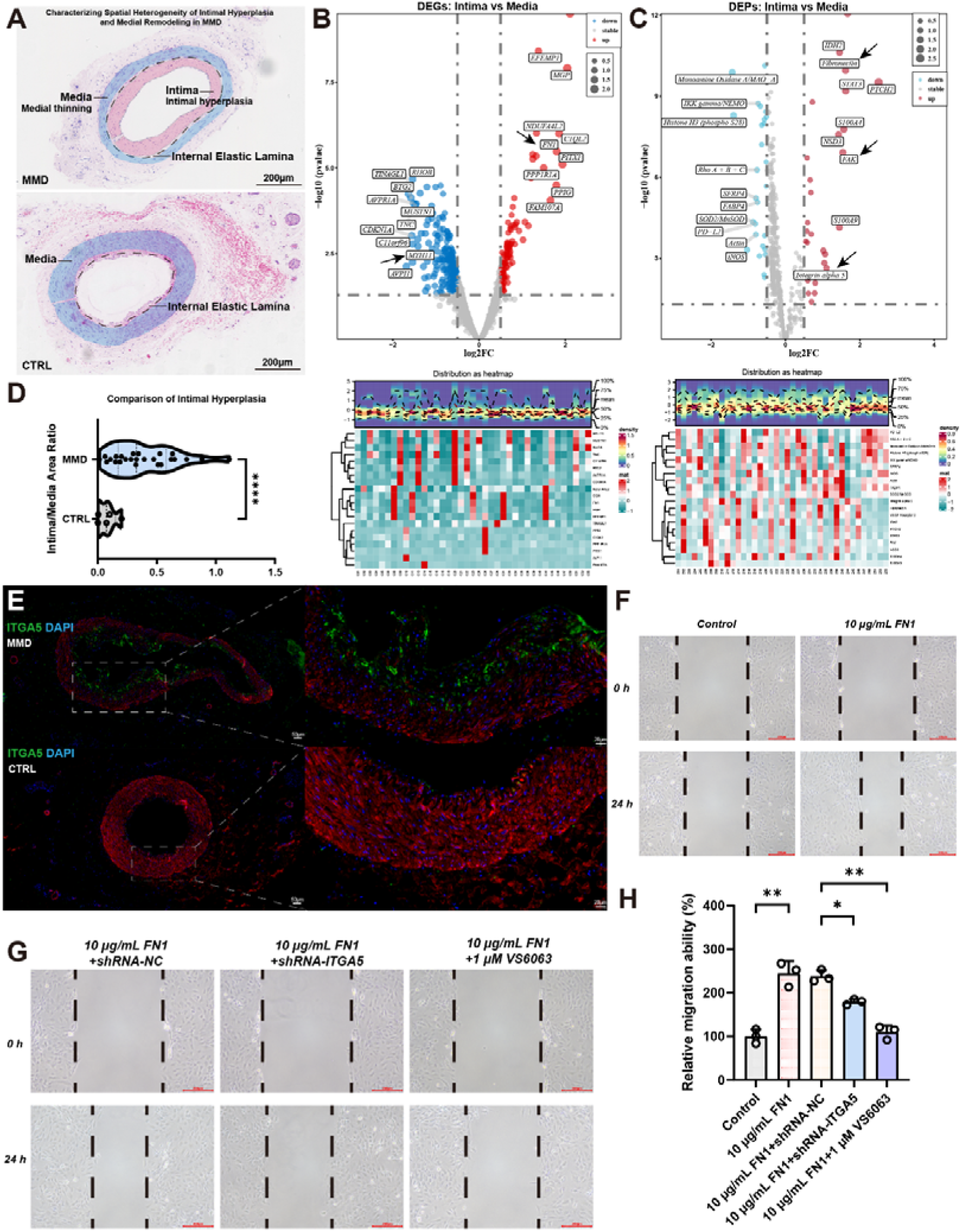
Spatial transcriptomic and proteomic profiling identifies the FN1-ITGA5-FAK axis enriched in the MMD intima. (A) Representative hematoxylin-eosin (HE) staining images of MMD and CTRL arteries, showing characteristic medial thinning and intimal hyperplasia, and based on this, a spatial omics ROI analysis was designed. (B-C) Volcano plots and corresponding heatmaps of differentially expressed genes (DEGs) and differentially expressed proteins (DEPs) between the intima and media of MMD arteries. *FN1*, *ITGA5*, and *FAK* are significantly upregulated in the intima. (D) Violin plot quantifying the Intima/Media Area Ratio in MMD versus CTRL groups (****p < 0.0001). (E) Representative IF staining of ITGA5 (green) demonstrating its robust accumulation in the thickened intima of MMD compared to CTRL. (F-H) Scratch wound-healing assay and quantification of relative migration ability in VSMCs treated with 10 μg/mL FN1, showing that FN1-induced hyper-migration is significantly abrogated by ITGA5 knockdown (shRNA-ITGA5) or FAK inhibition (VS6063).

Differential expression analysis identified a distinct transcriptional pattern in the hyperplastic intima, with increased expression of several genes related to extracellular matrix remodeling and vascular lesion formation, including FN1 and EFEMP1 together with reduced expression of the contractile smooth muscle marker MYH11. Heatmap visualization further described clear segregation of the intimal and medial transcriptomic profiles (Fig. 2B).

We then performed spatial proteomic analysis on the same regions. Consistent with the transcriptomic findings, the intimal compartment showed increased abundance of proteins involved in matrix deposition and cell adhesion, including fibronectin, integrin alpha 5, and FAK. The proteomic heatmap likewise showed distinct molecular differences between intima and media (Fig. 2C). Overall, these data show that the hyperplastic intima in MMD exhibits enhanced ECM and adhesion signaling together with loss of a contractile smooth muscle feature.

To validate the spatial profiling results at the tissue level, immunofluorescence staining was performed for ITGA5. Compared with control STA, MMD vessels showed stronger ITGA5 immunoreactivity, particularly in the thickened intimal region (Fig. 2E), supporting regional activation of the ITGA5-associated adhesion pathway in diseased arteries.

Given the enrichment of FN1 and ITGA5 in the intima, we next tested whether FN1 could directly enhance smooth muscle cell migration through the ITGA5–FAK axis. To investigate the functional role of ITGA5 in smooth muscle cells, we first evaluated three shRNA constructs targeting ITGA5 in HBVSMCs. shRNA-ITGA5#2 showed the strongest knockdown efficiency at both the protein and mRNA levels and was therefore used in subsequent experiments (Fig. S1A–C).

In HBVSMCs, treatment with 10 μg/mL FN1 significantly increased wound closure compared with untreated controls (Fig. 2F, H). This pro-migratory effect was not altered by control shRNA, but was markedly attenuated by ITGA5 knockdown or pharmacological inhibition of FAK with VS6063 (Fig. 2G, H). These findings support a model in which ECM remodeling in the MMD intima, particularly increased FN1, promotes smooth muscle cell migration through activation of the ITGA5–FAK pathway. Consistent with the migration results, western blot analysis showed that 10 μg/mL FN1 treatment reduced the expression of the contractile smooth muscle markers MYH11, CNN1, and TAGLN in HBVSMCs (Fig. S2A–D). In parallel, FN1 stimulation increased the expression of matrix-related proteins, including FN1, COL1A1, COL3A1, OGN, and EFEMP1 (Fig. S2A, E–I). These changes were not significantly altered by control shRNA, but were partially reversed by ITGA5 knockdown or FAK inhibition with VS6063, which restored contractile marker expression and reduced the levels of ECM-associated proteins (Fig. S2A–I). These data further support that FN1 promotes smooth muscle remodeling through the ITGA5–FAK pathway, accompanied by suppression of the contractile phenotype and induction of an ECM-associated molecular program. In parallel, we assessed activation of the downstream FAK pathway. Western blot analysis showed that FN1 treatment had no significant effect on total FAK expression, but significantly increased FAK phosphorylation at Tyr397 in HBVSMCs (Fig. S3A–C). This effect was unchanged in the shRNA-NC group, but was markedly attenuated by ITGA5 silencing or VS6063 treatment (Fig. S3A–C), further supporting activation of the FN1–ITGA5–FAK axis in smooth muscle cells.

### ITGA5 inhibition attenuates the migratory phenotype and FAK activation in MMD iPSC-derived SMCs

To further determine whether the ITGA5–FAK axis contributes to the abnormal phenotype of patient-derived smooth muscle cells, we generated iPSC-derived VSMCs from PBMCs of MMD and control subjects, followed by ITGA5 knockdown or pharmacological inhibition of FAK in MMD-iPSC-SMCs (Fig. 3A). The differentiated cells showed the expected α-SMA-positive smooth muscle phenotype in both the MMD and control groups, as confirmed by immunofluorescence staining (Figs. S4 and S5).

**Figure 3.**
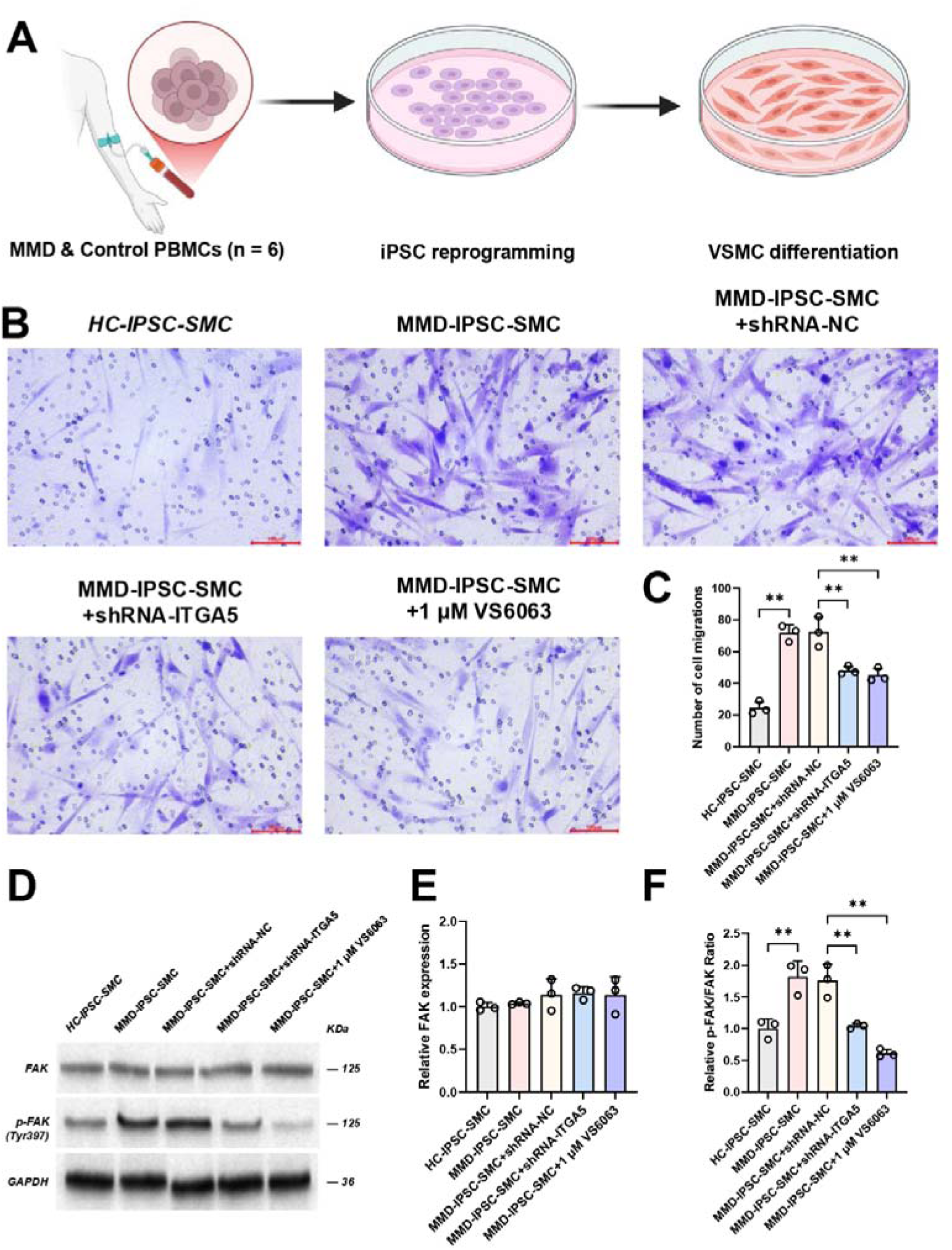
ITGA5-FAK signaling drives the hyper-migratory phenotype in MMD patient-derived iPSC-SMCs. (A) Workflow for the generation of induced pluripotent stem cells (iPSCs) from peripheral blood mononuclear cells (PBMCs) of MMD patients and healthy controls (HC), followed by directed differentiation into VSMCs. (B-C) Representative images and quantification of Transwell migration assays. MMD-iPSC-SMCs exhibit significantly enhanced migration compared to HC-iPSC-SMCs, which is effectively reversed by shRNA-ITGA5 or VS6063 treatment. (D-F) Western blot analysis and quantification of total FAK and phosphorylated FAK (p-FAK at Tyr397) in the established iPSC-SMC models. The elevated p-FAK/FAK ratio in MMD-iPSC-SMCs is substantially reduced by ITGA5 silencing or direct FAK inhibition.

Transwell assays showed that MMD-iPSC-SMCs displayed a markedly higher migratory capacity than HC-iPSC-SMCs (Fig. 3B,C). This increase was not altered by control shRNA, but was significantly reduced after ITGA5 silencing or treatment with 1 μM VS6063 (Fig. 3B,C), indicating that the enhanced migratory behavior of MMD-iPSC-SMCs is dependent, at least in part, on the ITGA5–FAK pathway.

We next examined downstream FAK signaling in these cells. Western blot analysis showed that total FAK expression was not significantly different among groups (Fig. 3D,E). In contrast, the p-FAK/FAK ratio was significantly increased in MMD-iPSC-SMCs compared with HC-iPSC-SMCs (Fig. 3D,F). This increase was not changed by shRNA-NC, but was markedly attenuated by shRNA-ITGA5 or VS6063 treatment (Fig. 3D,F).

Together, these results indicate that MMD iPSC-derived SMCs exhibit enhanced migration and increased FAK activation, and that both phenotypes can be attenuated by inhibition of the ITGA5–FAK axis (Fig. 3B–F).

Compared with HC-iPSC-SMCs, MMD-iPSC-SMCs showed lower expression of the contractile markers MYH11, CNN1, and TAGLN (Fig. 4A–D). In contrast, proteins associated with extracellular matrix remodeling, including FN1, COL1A1, COL3A1, OGN, EFEMP1, and ITGA5, were all increased (Fig. 4A, E–J). These changes were not appreciably altered by shRNA-NC. However, both ITGA5 silencing and FAK inhibition shifted the profile toward that of the control group, with recovery of MYH11, CNN1, and TAGLN and reduction of FN1, COL1A1, COL3A1, OGN, EFEMP1, and ITGA5 expression (Fig. 4A–J).

**Figure 4.**
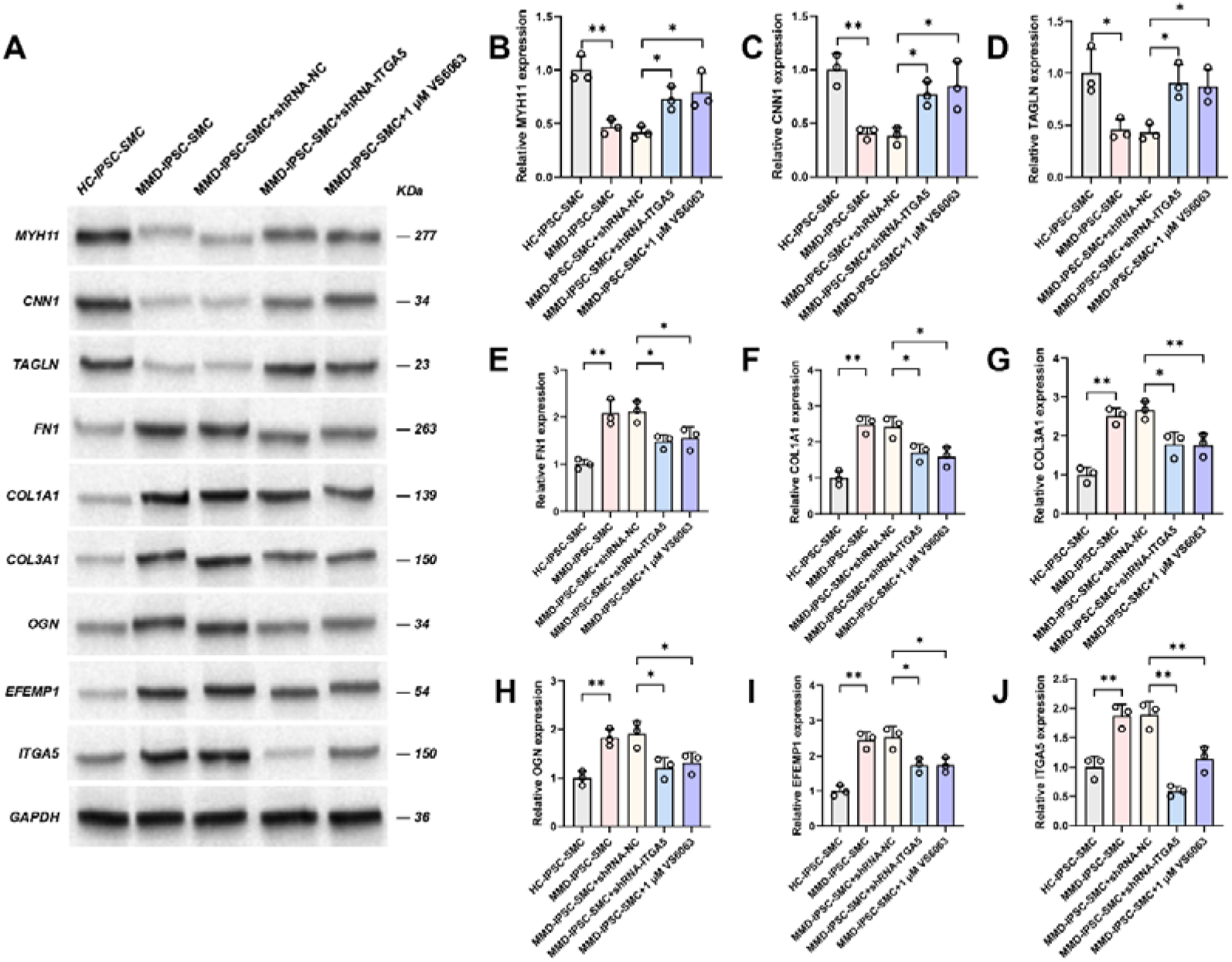
**Blocking the ITGA5 pathway may reduce the pathological phenotypic transformation of MMD-iPSC-SMC.**(A) Representative Western blot images showing the expression levels of SMC contractile markers, synthetic/matrix markers, and ITGA5 in HC-iPSC-SMCs and MMD-iPSC-SMCs under various treatments. (B-D) Quantification of contractile markers (MYH11, CNN1, TAGLN). Expression is significantly loss in MMD-iPSC-SMCs but rescued upon treatment with shRNA-ITGA5 or VS6063. (E-J) Quantification of synthetic and extracellular matrix-related proteins (FN1, COL1A1, COL3A1, OGN, EFEMP1) and ITGA5. The abnormal upregulation of these markers in MMD-iPSC-SMCs is suppressed by targeting the ITGA5-FAK axis, indicating a reversal from the synthetic back to the contractile phenotype.

Together, these findings show that MMD iPSC-derived SMCs display enhanced migration, increased FAK activation, loss of contractile markers, and upregulation of matrix-associated proteins. Interference with the ITGA5–FAK axis attenuated each of these abnormalities (Figs. 3B–F and 4A–J).

### Spatial metabolomics reveals distinct metabolic features in the hyperplastic intima in MMD

Spatial metabolomic profiling revealed marked regional heterogeneity across MMD arterial sections. Representative ion images showed that multiple metabolites displayed spatially restricted distributions within the vessel wall (Fig. 5A), and 3D UMAP analysis further supported the existence of distinct metabolomic states across tissue regions (Fig. 5B).

**Figure 5.**
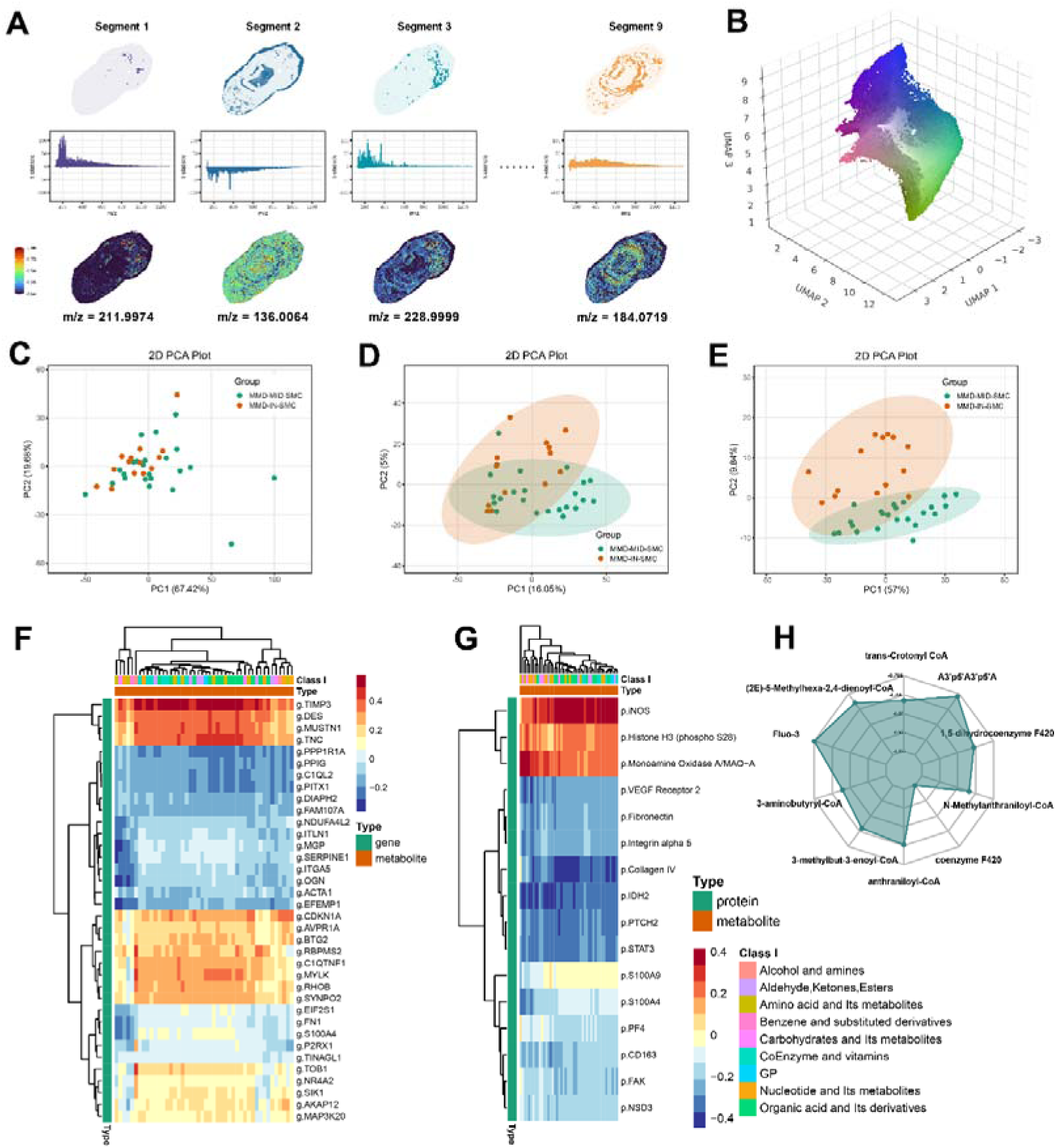
Integrated spatial multi-omics reveals metabolic reprogramming associated with the FN1-ITGA5 axis in MMD SMCs. (A) Representative spatial metabolomics mapping (m/z signals) showing the heterogeneous distribution of specific metabolites across different arterial segments. (B) 3D UMAP projection of spatial metabolic segments. (C-E) 2D Principal Component Analysis (PCA) plots distinguishing the distinct molecular signatures between MMD-MID-SMC (media) and MMD-IN-SMC (intima). (F-G) Multi-omics correlation heatmaps illustrating the strong spatial co-expression and inverse correlations between specific intimal genes/proteins (e.g., *g.ITGA5*, *p.Fibronectin*) and the abundance of localized metabolites. (H) Radar chart highlighting the top depleted compounds in the MMD intima compared to the media, revealing a profound exhaustion of Acyl-CoA derivatives (e.g., trans-Crotonyl CoA, 3-aminobutyryl-CoA).

We then compared metabolomic signatures between intimal and medial smooth muscle cell-rich areas. PCA found separation between MMD-IND-SMC and MMD-MID-SMC groups, indicating metabolic differences between the two compartments (Fig. 5C–E).

### Integrated multi-omics correlation analysis links intimal remodeling to a distinct metabolically depleted niche

To further define how metabolic alterations were coupled to spatial transcriptional and proteomic changes, we performed integrated correlation analyses between metabolites and region-resolved transcriptomic and proteomic features. In the joint transcriptome–metabolome heatmap, hierarchical clustering separated genes into distinct modules with sharply different metabolic correlation patterns (Fig. 5F). One prominent cluster was composed of genes associated with intimal remodeling and synthetic smooth muscle features, including ITGA5, OGN, EFEMP1, and SERPINE1. These genes were concentrated within a broad negative-correlation block, indicating that their increased expression was accompanied by reduced abundance of a large set of metabolites (Fig. 5F). This pattern suggests that the ITGA5-dominant intimal remodeling program is embedded within a metabolically altered microenvironment characterized by relative metabolite depletion, consistent with increased metabolic consumption or rewiring during smooth muscle migration and matrix production.

In contrast, a separate gene module showed an opposite association pattern. Genes such as DES and TIMP3, which are more consistent with a stable or contractile smooth muscle state, clustered within a positive-correlation region (Fig. 5F). This result indicates that the medial contractile program is linked to a different metabolic context from that of the hyperplastic intima. Thus, the transcriptome–metabolome correlation structure identifies a clear metabolic boundary between contractile medial SMCs and remodeling-associated intimal SMCs.

Notably, FN1 did not cluster tightly with the main ITGA5/OGN/EFEMP1 negative-correlation module. Instead, it localized to a nearby but distinct branch together with S100A4 (Fig. 5F). This distribution suggests that, within the spatially complex lesion microenvironment, FN1 transcription may not fully mirror ITGA5 expression dynamics.

Rather, FN1 may be more closely linked to local injury or inflammatory remodeling signals, whereas ITGA5 upregulation may reflect the cellular response phase that executes matrix sensing, adhesion, and migration.

We next performed proteome–metabolome correlation analysis. This heatmap also resolved several biologically distinct clusters (Fig. 5G). At the top, iNOS, phospho-Histone H3 (S28), and MAO-A formed a positive-correlation module, indicating that inflammatory or stress-associated protein expression was linked to relative enrichment of multiple metabolites (Fig. 5G). In the middle, a second and highly relevant module contained VEGF Receptor 2, fibronectin, integrin alpha 5, and Collagen IV, all of which showed broad negative correlations with the metabolite panel (Fig. 5G). Importantly, fibronectin and integrin alpha 5 were positioned very closely in the hierarchical tree, supporting strong spatial and biological coupling of this ligand–receptor pair within the lesion microenvironment. The overall negative-correlation pattern of this module suggests that regions enriched for ECM deposition, adhesion signaling, and vascular remodeling exist in a relatively metabolite-poor state.

A third module, including S100A9, S100A4, CD163, and FAK, showed weaker and more mixed correlations with metabolites (Fig. 5G). Unlike fibronectin and integrin alpha 5, FAK did not cluster within the tight ECM-associated branch, suggesting that its spatial distribution may reflect additional regulation by the local inflammatory microenvironment rather than by matrix accumulation alone. Together, these data reveal two major metabolic ecologies within the MMD vessel wall: an inflammation-associated, metabolically enriched niche and an FN1/ITGA5-centered matrix-remodeling niche associated with relative metabolite depletion.

To identify the metabolites most strongly altered between intimal and medial SMC-rich regions, we further summarized the top differential metabolites by fold change. The radar plot showed that the 10 most altered metabolites were uniformly decreased in intimal SMCs compared with medial SMCs (Fig. 5H). These included several acyl-CoA–related intermediates and metabolites linked to coenzyme/vitamin metabolism, amino acid-related pathways, and small-molecule biosynthesis (Fig. 5H). The consistent downward shift of these metabolites indicates that the hyperplastic intima is associated with broad depletion of key metabolic intermediates rather than selective accumulation of a few species.

## Discussion

To our knowledge, this is the first study to integrate multi-sample STA spatial transcriptomics, spatial proteomics, spatial metabolomics, single-cell analysis, and iPSC-SMC functional validation in one MMD framework. Our data indicate that the hyperplastic intima in MMD is not merely a morphologic thickening, but a lesion with a definable molecular identity that includes extracellular matrix remodeling, activation of the ITGA5–FAK axis, loss of contractile smooth muscle markers, and a metabolically depleted niche. The consistency between spatial data and the two functional systems, namely FN1-stimulated HBVSMCs and patient-derived iPSC-SMCs, strengthens the biological coherence of this model. Taken together, these findings provide a spatially resolved mechanistic framework for intimal hyperplasia in MMD.

Classic histopathological studies of MMD have consistently documented hallmark vascular lesions, including concentric intimal hyperplasia, medial thinning, and fragmentation of the internal elastic lamina.^18^ Our Hematoxylin and Eosin (H&E) and alpha smooth muscle actin staining results align with these classical observations. Further analysis of these results reveals that the thickened intima represents an arterial remodeling process, accompanied by a large accumulation of α-SMA-positive cells. This pattern raises the possibility that smooth muscle related cells leave their original medial program and enter a state dominated by migration, growth, and matrix production. However, traditional morphological evaluations inherently fail to explain why the intimal zone becomes the dominant pathological site in MMD.^19^ While recent bulk RNA sequencing and single cell transcriptomics have identified potential cellular perturbations in MMD, these techniques inevitably lose crucial spatial context.^20^ Several recent studies have already pointed toward molecular abnormalities in MMD arteries, but each captured only one layer of the disease. Bulk transcriptomics of intracranial arteries highlighted ECM and mitochondrial pathways as major signals, yet by design could not resolve which arterial compartment carried those programs.^21^ Single-cell analysis of STA tissue then showed that smooth muscle and immune populations are altered in MMD, but dissociation removes the physical relationship between cell states and lesion topography.^14^ In recent literature, several ECM related studies in MMD have highlighted similar trends. For instance, Wang et al. provided evidence that altered MMP-9 interacting with Desmoglein-2 accelerates intimal thickening, which aligns with our observed *SERPINE1* and matrix dysregulation.^22^ Similarly, Xu et al. reported elevated collagen and matrix degradation factors in MMD middle cerebral arteries, further supporting our finding that a continuous turnover of the ECM scaffold provides the physical space for intimal expansion.^21^ Notably, their bulk transcriptomic analysis also highlighted oxidative phosphorylation and mitochondrial dysfunction alongside ECM organization as top dysregulated pathways, strongly echoing our hypothesis of coupled metabolic-structural rewiring. In this Study, our spatial transcriptomics revealed a pronounced spatial enrichment of ECM related genes, such as FN1 and EFEMP1, specifically within the hyperplastic intima. Correspondingly, our spatial proteomics confirmed the significant upregulation of ITGA5, the classical receptor for fibronectin. Although the exact spatial dynamics of this axis in MMD remains elusive, extensive studies in other occlusive vascular diseases, such as atherosclerosis and restenosis after angioplasty, have clearly suggested that the FN1-ITGA5 interaction is a major regulator of vascular smooth muscle cell (VSMC) plasticity.^23^ Upon binding to the provisional FN1 matrix, ITGA5 initiates robust “outside-in” signaling cascades, predominantly through the activation of focal adhesion kinase (FAK), which directly orchestrates the phenotypic switching of VSMCs from a quiescent, contractile state to a highly migratory, synthetic phenotype.^24,25^ Consistent with the mechanism, our in vitro FN1 stimulation experiments suggest that this fibronectin-rich environment can effectively drive cell migration, suggesting that this may be a key signaling pathway for endometrial hyperplasia in Moyamoya disease.

A central consequence of this ECM remodeling is the phenotypic transition of SMCs. Historically, *MYH11*, *CNN1*, and *TAGLN* are recognized as classical markers of the fully differentiated contractile SMC state.^26^ Our study is spatially map the profound downregulation of *MYH11* strictly within the MMD intimal region, representing an explicit spatial suppression of the contractile program. Recent advances in MMD research have further underscored the complexity of this process.^27^ For instance, a recent spatial proteomics analysis corroborated that increased intimal fibronectin deposition directly mirrors the contractile-to-synthetic phenotypic switching of VSMCs in MMD arteriopathy.^28^

Mechanistically, this loss of contractile identity is frequently governed by master transcription factors such as Krüppel-like factor 4 (KLF4), which acts as a pivotal switch to repress VSMC differentiation genes and promote a highly migratory state.^29^ In the specific context of MMD, our previous organoid and single-cell profiling models observed that serum-derived factors, such as specific tubulin proteins, can trigger this transition by activating the PI3K/AKT/KLF4 signaling cascade, ultimately driving VSMC migration and intimal accumulation.^13^ Additionally, studies indicating that VEGF and angiopoietin imbalances induce abnormal matrix remodeling in MMD are highly consistent with our spatial mapping of VEGFR2 and FN1 co localization.^30^ Because standard transgenic murine models currently fail to fully recapitulate the spontaneous intimal hyperplasia of human MMD.^31^ Our patient-derived, specific iPSC-SMC model provides some functional relevance to in vitro studies. Through this platform, we established that the ITGA5–FAK axis is not merely a spatial co localization marker, but rather a functional mediator possessing targetable properties.^32^ This active mediatory role positions the ITGA5–FAK signaling cascade as a highly promising candidate for further mechanistic intervention.^29,33^

The intimal SMC rich region is characterized by a global depletion of various metabolites, with acyl-CoA related metabolites ranking among the most drastically reduced compounds. Acyl-CoA derivatives are central to energy production and lipid metabolism, and disturbance or depletion of acyl-CoA–related intermediates has been linked to vascular metabolic stress and dysfunction.^34,35^ In our transcriptome metabolome correlation heatmap, highly expressed intimal genes (*ITGA5*, *OGN*, *EFEMP1*, *SERPINE1*) fell into a broad negative correlation zone with these metabolites. This indicates that ECM remodeling is not an isolated event; rather, it occurs within a pathologically active niche where essential metabolites are relatively deficient or rapidly consumed.^36^ We suggest that SMCs residing in the hyperplastic intima may be undergoing intense metabolic reprogramming. In a recent serum metabolomics study, we identified marked metabolic abnormalities across MMD subgroups and highlighted reduced Lysophosphatidylcholine 16:1-2, indicating that abnormal vascular remodeling in MMD is accompanied by a measurable metabolic disturbance rather than being a purely structural process.^37^ The concept of metabolic microenvironment as a driver of intimal hyperplasia is further supported by recent investigations into other stenotic vascular diseases. For instance, Zhao et al. described that venous intimal hyperplasia in arteriovenous fistulas is characterized by a broad spectrum of metabolic dysregulation, particularly involving amino acid and lipid metabolism pathways, which directly correlates with accelerated VSMC proliferation and pathological remodeling.^38^ Given the central role of acyl-CoA intermediates in fatty acid handling, acetyl group transfer, and cellular bioenergetics, depletion of this pool may reflect increased metabolic demand during migration and matrix remodeling.^39^ You et al. reported that mitochondrial Ac-CoA accumulation and its cytoplasmic transfer regulate histone acetylation at the KLF4 promoter and thereby influence VSMC phenotypic switching, which suggests that disruption of acyl-CoA homeostasis can directly intersect with the transcriptional machinery that drives smooth muscle remodeling; in our dataset, the marked loss of acyl-CoA–related metabolites in intimal SMC-rich regions may therefore reflect a similarly reprogrammed state, although the precise flux direction remains to be determined.^40^

## Conclusions

Currently, the clinical management of MMD remains heavily reliant on surgical revascularization^17^, largely due to the critical lack of mechanistic pharmacological agents. Although the superficial temporal artery (STA) does not represent the terminal intracranial ICA, its histological consistency and surgical accessibility render it an invaluable surrogate window for investigating MMD vascular remodeling. Our findings suggest that the ITGA5–FAK signaling axis may serve as a potential therapeutic entry point, a concept that warrants continued validation across patient derived organoids and suitable animal surrogates.

## Limitations

Several limitations deserve note. First, although the STA provides a practical and informative surrogate tissue, it does not fully recapitulate the hemodynamic and molecular features of the terminal intracranial lesions in MMD. The control cohort was modest, and the spatial transcriptomic, proteomic, and metabolomic arms were necessarily limited by tissue availability, so external validation in larger series will be important. Regarding the metabolomic landscape, although the marked localized metabolite decline incontrovertibly signifies metabolic rewiring and depletion, the static nature of spatial mass spectrometry precludes definitive assertions concerning directional metabolic fluxes; thus, orthogonal validation via isotope tracing or targeted metabolic assays remains an indispensable future endeavor. Ultimately, while the iPSC-SMC model offers a useful human platform for mechanistic studies, it cannot fully reproduce the complex in vivo vascular environment, including shear stress, inflammatory infiltration, and long term biomechanical remodeling.

## Acknowledgement

We thank the healthcare staff in the department for their assistance with this project. We also sincerely thank all participants for their support and cooperation.

## Funding Sources

This study was supported by Natural Science Foundation of China (grant no. 82471337;82471328; 82371296).

## Authors’ contributions

Shihao He, Xun Ye, Rong Wang, and Yuanli Zhao conceived and designed the study. Shihao He, Xiaofan Yu, Yaoren Chang, Zhenyu Zhou, and Junze Zhang collected clinical samples and performed clinical data entry. Shihao He, Xiaofan Yu, Talha Ahmed, Yifan Xu, Hanzhi Liu, Shaoqi Xu, and Lucas Du conducted spatial multi-omics analysis, bioinformatic analysis, and data visualization. Shihao He led the data integration, interpreted the results, and drafted the manuscript. Xun Ye, Rong Wang, and Yuanli Zhao revised the manuscript critically for important intellectual content. All authors reviewed and approved the final manuscript.

## Competing interests

The authors declare no competing interests.

## Consent for publication

Not applicable.

## Data availability statement

The data presented in the current study are available from the corresponding author upon reasonable request.

## Notes

### Competing Interest Statement

The authors have declared no competing interest.

