## Supplementary Table1 for "Spatial Multi-Omics Reveals Extracellular Matrix Remodeling and VSMC Phenotypic Switching in Moyamoya Disease"

**Supplementary Materials**

**Table S1. Primers used in Quantitative Real-Time PCR**

| Primers | Sequence (5'→3') | |
|---|---|---|
| ITGA5 | Forward | CGGGCTCCTTCTTCGGATTC |
| | Reverse | ATTCAATGGGGGTGCACTGT |
| GAPDH | Forward | AATGGGCAGCCGTTAGGAAA |
| | Reverse | GCGCCCAATACGACCAAATC |

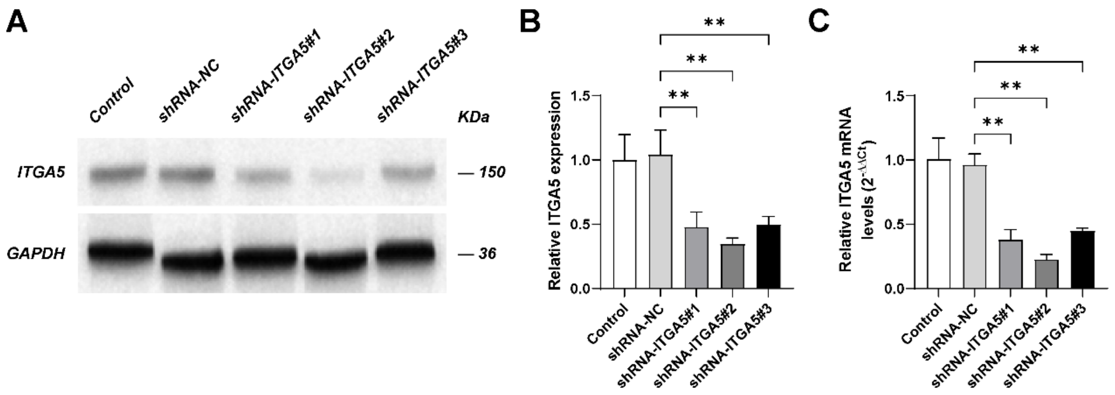


**Figure S1. Screening of shRNA constructs targeting ITGA5 in HBVSMCs.**

(A) Western blot analysis of ITGA5 expression after transduction with shRNA-NC or three ITGA5-targeting shRNAs. GAPDH served as the loading control.

(B) Quantification of ITGA5 protein levels.

(C) qPCR validation of ITGA5 mRNA knockdown efficiency. Data are shown as mean ± SD from three independent experiments. $P < 0.01$ by one-way ANOVA with Tukey's post hoc test.

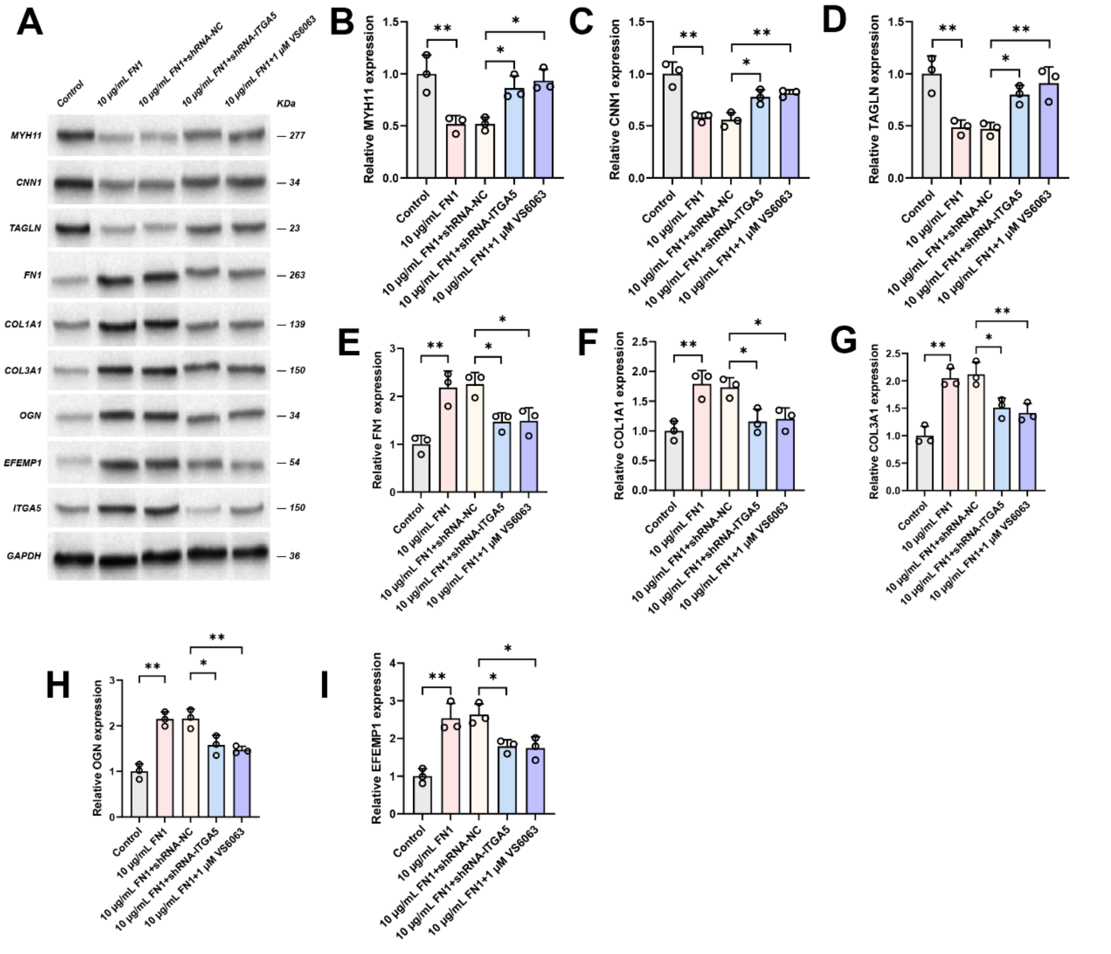


**Figure S2. ITGA5 knockdown or FAK inhibition attenuates FN1-induced protein changes in HBVSMCs.**

(A) Representative western blots showing the expression of MYH11, CNN1, TAGLN, FN1, COL1A1, COL3A1, OGN, EFEMP1, and ITGA5 in HBVSMCs under the indicated conditions: control, 10 µg/mL FN1, 10 µg/mL FN1 + shRNA-NC, 10 µg/mL FN1 + shRNA-ITGA5, and 10 µg/mL FN1 + 1 µM VS6063. GAPDH was used as the loading control.

(B–D) Quantification of the contractile smooth muscle markers MYH11, CNN1, and TAGLN.

(E–I) Quantification of FN1, COL1A1, COL3A1, OGN, and EFEMP1 expression.

Data are presented as mean ± SD from three independent experiments. Statistical significance was determined by one-way ANOVA followed by Tukey's multiple-comparisons test. *P* < 0.05, P < 0.01.

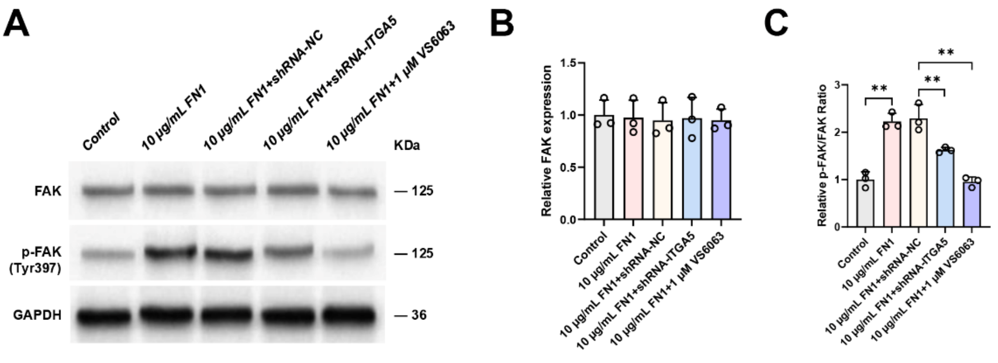


**Figure S3. Effects of ITGA5 knockdown and FAK inhibition on FN1-induced FAK phosphorylation in HBVSMCs.**

(A) Representative western blots showing total FAK and phosphorylated FAK (Tyr397) levels in HBVSMCs under the indicated conditions: control, 10 µg/mL FN1, 10 µg/mL FN1 + shRNA-NC, 10 µg/mL FN1 + shRNA-ITGA5, and 10 µg/mL FN1 + 1 µM VS6063. GAPDH was used as the loading control.

(B) Quantification of total FAK protein expression.

(C) Quantification of the p-FAK/FAK ratio.

Data are presented as mean ± SD from three independent experiments. Statistical significance was determined by one-way ANOVA followed by Tukey's multiple-comparisons test. *P* < 0.05, **P** < 0.01.

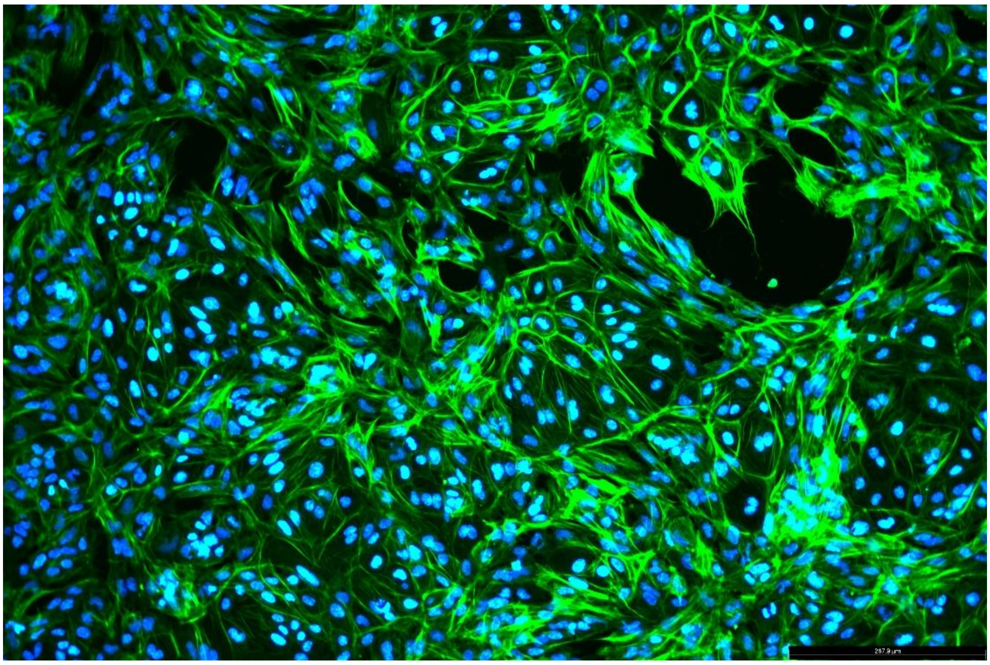

**Figure S4. Representative α-SMA immunofluorescence staining of MMD iPSC-derived VSMCs.**

Differentiated MMD iPSC-derived VSMCs were stained with α-SMA (green) and counterstained with DAPI (blue).

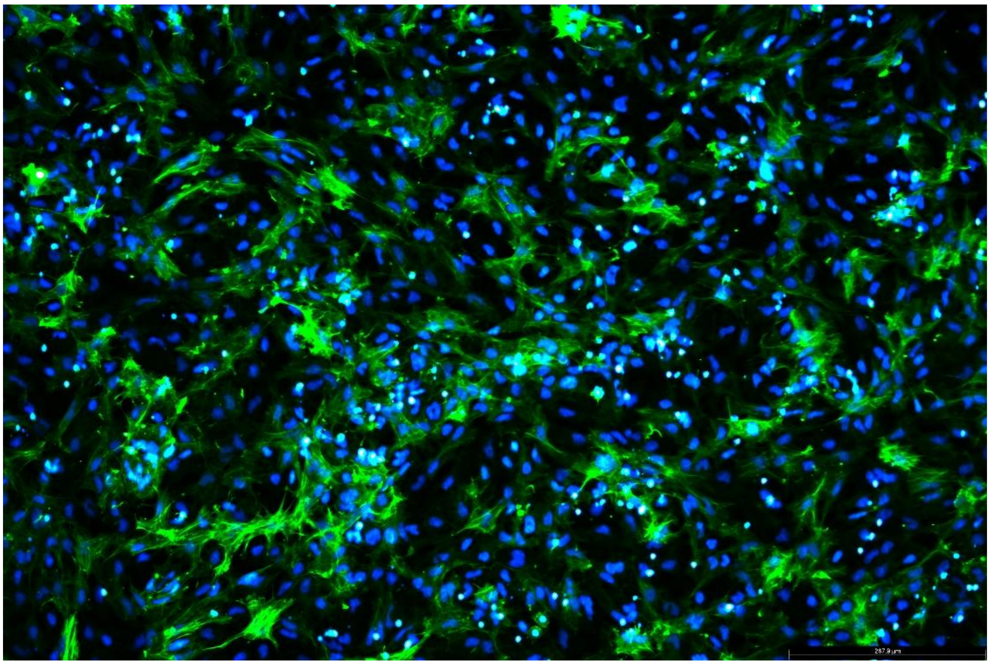

**Figure S5. Representative α-SMA immunofluorescence staining of Control iPSC-derived VSMCs.**

Differentiated Control iPSC-derived VSMCs were stained with α-SMA (green) and counterstained with DAPI (blue).
